# Drosophila embryonic type II neuroblasts: origin, temporal patterning, and contribution to the adult central complex

**DOI:** 10.1101/170043

**Authors:** Kathleen T. Walsh, Chris Q. Doe

## Abstract

Drosophila neuroblasts are an excellent model for investigating how neuronal diversity is generated. Most brain neuroblasts generate a series of ganglion mother cells (GMCs) that each make two neurons (type I lineage), but sixteen brain neuroblasts generate a series of intermediate neural progenitors (INPs) that each produce 4-6 GMCs and 8-12 neurons (type II lineage). Thus, type II lineages are similar to primate cortical lineages, and may serve as models for understanding cortical expansion. Yet the origin of type II neuroblasts remains mysterious: do they form in the embryo or larva? If they form in the embryo, do their progeny populate the adult central complex, as do the larval type II neuroblast progeny? Here we present molecular and clonal data showing that all type II neuroblasts form in the embryo, produce INPs, and express known temporal transcription factors. Embryonic type II neuroblasts and INPs undergo quiescence, and produce embryonic-born progeny that contribute to the adult central complex. Our results provide a foundation for investigating the development of the central complex, and tools for characterizing early-born neurons in central complex function.

## INTRODUCTION

Drosophila neural progenitors, called neuroblasts, are a model system for investigating stem cell self-renewal versus differentiation (Doe, 2008; Reichert, 2011), as well as how a single progenitor generates different types of neurons and glia over time (Alsio et al., 2013; Kohwi et al., 2013). Drosophila type I neuroblasts have a relatively simple cell lineage: they undergo a series of asymmetric cell divisions to produce a series of smaller ganglion mother cells (GMCs) that typically differentiate into a pair of neurons. There are about 100 type I neuroblasts in each larval brain lobe; they generate progeny during embryogenesis, undergo a period of quiescence, and then resume their lineage in the larva (Truman and Bate, 1988; Datta, 1995; Maurange and Gould, 2005; Sousa-Nunes et al., 2010). Type I neuroblasts have a molecular profile that is Deadpan (Dpn)^+^, Asense (Ase)^+^ and Pointed P1 (PntP1)^-^ (Zhu et al., 2011; Xie et al., 2016). Moreover, many embryonic type I neuroblasts can transition to a simpler “type 0” lineage, in which each neuroblast daughter cell directly differentiates into a neuron (Karcavich and Doe, 2005; Baumgardt et al., 2014; Bertet et al., 2014). In contrast, Drosophila type II neuroblasts have a more elaborate cell lineage: they divide asymmetrically to bud off smaller intermediate neural progenitors (INPs) that themselves produce a series of 4-6 GMCs that each make a pair of neurons or glia (Bello et al., 2008; Boone and Doe, 2008; Bowman et al., 2008; Izergina et al., 2009). Type II neuroblasts have a molecular profile that is Dpn^+^Ase^-^ PntP1^+^ (Bello et al., 2008; Boone and Doe, 2008; Bowman et al., 2008; Izergina et al., 2009; Zhu et al., 2011). Although there are only eight type II neuroblasts per larval brain lobe, they generate a major portion of the intrinsic neurons of the adult central complex (Bayraktar et al., 2010; Ito et al., 2013; Riebli et al., 2013; Yu et al., 2013), a neuropil devoted to multimodal sensory processing and locomotion (Martin et al., 1999; Renn et al., 1999; Strauss, 2002; Wessnitzer and Webb, 2006; Poeck et al., 2008; Wang et al., 2008; Pan et al., 2009; Bender et al., 2010; Boyan and Reichert, 2011; Ofstad et al., 2011; Seelig and Jayaraman, 2011; Seelig and Jayaraman, 2013; Seelig and Jayaraman, 2015).

A large amount of work over the past two decades has illuminated the general principles for how type I neuroblasts generate neuronal diversity. First, dorso-ventral, anterior-posterior, and Hox spatial patterning cues generate unique neuroblast identities (Chu-LaGraff and Doe, 1993; Prokop and Technau, 1994; Skeath et al., 1995; McDonald et al., 1998; Weiss et al., 1998; Skeath and Thor, 2003; Marin et al., 2012; Estacio-Gomez and Diaz-Benjumea, 2014; Moris-Sanz et al., 2015). Second, the temporal transcription factors Hunchback (Hb), Krüppel (Kr), Nubbin/Pdm2 (Pdm), Castor (Cas) and Grainy head (Grh) specify unique GMC identities within each neuroblast lineage (Brody and Odenwald, 2000; Berger et al., 2001; Isshiki et al., 2001; Novotny et al., 2002; Cenci and Gould, 2005; Kanai et al., 2005; Grosskortenhaus et al., 2006; Mettler et al., 2006; Urban and Mettler, 2006; Maurange et al., 2008; Tran and Doe, 2008; Tsuji et al., 2008; Ulvklo et al., 2012; Herrero et al., 2014; Moris-Sanz et al., 2014). In contrast, much less is known about type II neuroblasts. Only one of the eight type II neuroblasts has been identified in the embryo (Hwang and Rulifson, 2011); the origin of the other type II neuroblasts has not been reported in existing embryonic brain neuroblast maps (Urbach and Technau, 2003). It remains unknown whether type II neuroblasts arise de novo from the neuroectoderm similar to type I neuroblasts, or whether they arise from a type I > type II transition similar to the type I > type 0 neuroblast transitions (Baumgardt et al., 2014; Bertet et al., 2014). If type II neuroblasts form during embryogenesis, it is unknown whether they utilize the same Hb > Kr > Pdm > Cas > Grh temporal transcription factor cascade to generate neuronal diversity, or whether they make embryonic born INPs that sequentially express Dichaete (D) > Grh > Eyeless similar to larval INPs (Bayraktar and Doe, 2013). Furthermore, if type II neuroblast lineages are initiated in the embryo, it would be interesting to know if their INPs undergo quiescence, similar to type I and II neuroblasts; if so they would be the only cell type beyond neuroblasts known to enter quiescence at the embryo/larval transition. Perhaps most importantly, identifying embryonic type II neuroblasts is essential for subsequent characterization of their early-born progeny, which are likely to generate pioneer neurons crucially important for establishing larval or adult brain architecture.

Here we address all of these open questions. We show that all eight type II neuroblasts form during embryogenesis. We use molecular markers and clonal data to show that embryonic type II neuroblasts give rise to INPs that produce multiple GMCs and neurons during embryogenesis, and that INPs undergo quiescence during the embryo-larval transition. We find that embryonic type II neuroblasts sequentially express a subset of neuroblast temporal transcription factors (Pdm > Cas > Grh), and embryonic INPs express a subset of the known larval INP temporal transcription factors (Dichaete). Finally, we show that embryonic INPs give rise to neurons that survive to populate the adult central complex.

## RESULTS

### All type II neuroblasts arise during embryogenesis

Larval type II neuroblasts are PntP1^+^ Dpn^+^ Ase^-^ and here we used these markers to determine if type II neuroblasts exist in the embryo. We found that type II neuroblasts formed internal to the dorsal cephalic neuroectoderm beginning at late stage 11. At this stage, there is one PntP 1^+^ Dpn^+^ Ase^-^ type II neuroblast in a stereotyped dorsal posteromedial location; this is always the first type II neuroblast to appear (Fig. 1). By stage 12, the number of type II neuroblasts along the dorso-medial region of the brain increased from four (8h) to six (9.5h), and from stage 15 (12h) to the end of embryogenesis there were reliably eight type II neuroblasts per lobe (Fig. 1), the same number previously observed at all stages of larval development (Bello et al., 2008; Boone and Doe, 2008; Bowman et al., 2008; Izergina et al., 2009). We reliably found three clusters of type II neuroblasts: an anteromedial group of three neuroblasts, a medial group of three neuroblasts, and a posterior ventrolateral group of two neuroblasts (Fig. 1A; summarized in Fig. 1B). Due to the dynamic morphogenetic movements of head involution, and the close positioning of the type II neuroblasts, we could not reliably identify individual neuroblasts within each cluster.

**Figure 1.**
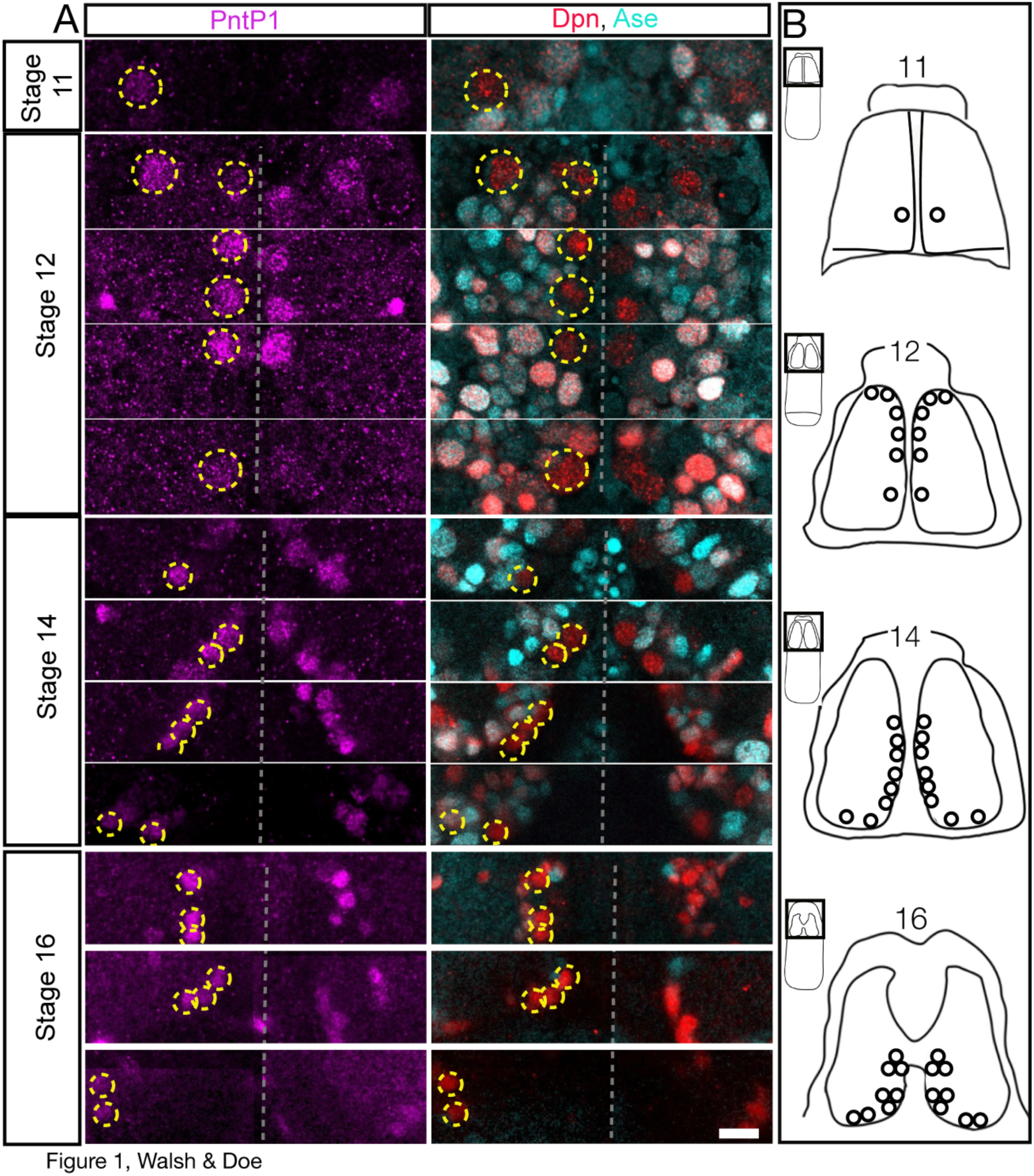
Eight type II neuroblasts arise during embryogenesis. **(A)** Embryonic type II neuroblasts (yellow circles on left brain lobe; unlabeled on right brain lobe) are PntP1+ (magenta) Dpn+ (red) Ase-(cyan)., Each stage shows multiple focal planes from anterior to posterior (top to bottom in the figure) to clearly visualize each type II neuroblast, except for stage 11 where there is a single type II neuroblast. **(B)** Summary of type II neuroblast formation; due to rapid morphogenetic movements it is not possible to identify individual type II neuroblasts from stage to stage, but beginning at stage 14 it is possible to recognize three clusters of neuroblasts. All panels are dorsal views with the dorsal midline in the center of the panel, anterior up. Scale bar = 10 μm.

We tried to link the embryonic type II neuroblasts to the map of embryonic brain neuroblasts (Urbach and Technau, 2003), but were unsuccessful, probably because most type II neuroblasts arise later than the stages described in that study. Based on molecular marker analysis, we conclude that all eight known type II neuroblasts form during embryogenesis and they are among the last neuroblasts to form during embryogenesis.

### Embryonic type II neuroblasts generate INPs, GMCs, and neurons during embryogenesis

Here we use molecular markers and clonal analysis to determine whether embryonic type II lineages produce INPs, GMCs, and neurons. We used a *Pnt-gal4* line to make clones; to validate the type II lineage-specific expression of this line, we stained for Pnt-gal4 and type II neuroblast and INP markers (Fig. 2A). We found that Pnt-gal4 is expressed in the parental type II neuroblast, the maturing INPs, and their GMC progeny (Fig. 2B). We did not detect any type I neuroblasts expressing this marker. Next, we generated “flip-out” clones using the heat shock-inducible multicolor flip out method (Nern et al., 2015) crossed to the *Pnt-gal4* line. When we assayed clones relatively early in embryogenesis (stage 13) we detected small clones containing a single type II neuroblast and one or more INPs (Fig. 2C; Table 1). Allowing the embryos to develop further resulted in larger clones that additionally contained GMCs and neurons (Fig. 2D). We found clones containing one type II neuroblast with up to five INPs at the latest stages of embryogenesis (Table 1). Taken together, these data show that embryonic type II neuroblasts generate multiple INPs which themselves produce GMCs and neurons prior to larval hatching.

**Figure 2.**
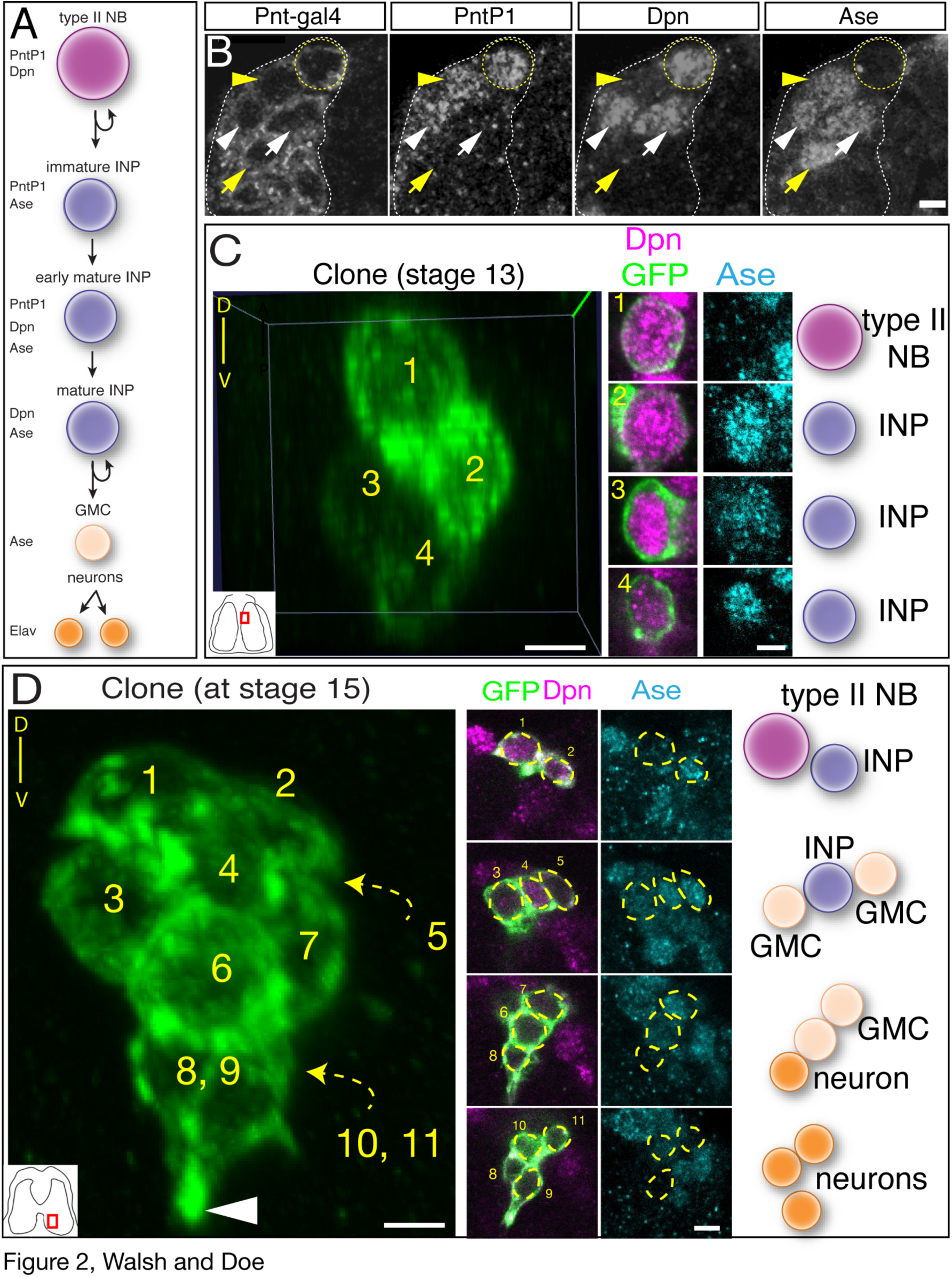
Clonal analysis shows that type II neuroblasts make INPs, GMCs and neurons during embryogenesis. **(A)** Molecular markers used to identify cell types within type II lineages, neuroblast (NB). **(B)** Embryonic type II neuroblasts generate embryonic-born INPs and GMCs. Dorso-medial view of a type II neuroblast cluster in a stage 16 embryo. Type II neuroblast (Pnt-gal4^+^ PntP1^+^ Dpn^+^ and Ase^-^; yellow circle); immature INP (Pnt-gal4^+^ PntP1^+^ Dpn^-^ and Ase^+^; yellow arrowhead); mature INP (Pnt-gal4^+^ PntP1^+^ Dpn^+^ and Ase^+^; white arrowhead); mature INP that has lost PntP1 expression (Pnt-gal4^+^ PntP1^-^ Dpn^+^ and Ase^+^; white arrow); and GMC (Pnt-gal4^+^ PntP1^-^ Dpn^-^ and Ase^+^; yellow arrow). Scale bar, 5 μm. **(C)** Single neuroblast clone assayed at stage 13; location shown in inset, lower left. Four cell clone containing a type II neuroblast and three INPs. Orientation is dorsal up, with the neuroblast closest to the dorsal surface of the brain. **(D)** Single neuroblast clone assayed at stage 15; location shown in inset, lower left. Eleven cell clone containing a type II neuroblast, two INP, four GMCs, and four neurons. Orientation is dorsal up, showing that the neurons are sending projections ventrally (arrowhead). Scale bar for (C) and (D) = 10 μm for clone projection, 5 μm for insets.

**Table 1.** Type II neuroblast clones contain INPs, GMCs, and neurons. Each row represents a single clone that is clearly spatially separate from other clones in the embryonic brain. Stage, time of clone analysis. Markers, molecular marker profile of each cell in the clone.

| Cluster | Stage | Type II<br>NB<br>*Dpn+ Ase-* | INP<br>*Dpn+Ase+* | GMC<br>*Dpn-Ase+* | Neuron<br>*Dpn- Ase-* | Total<br>Cells |
| --- | --- | --- | --- | --- | --- | --- |
| Anterior | 15 | 1 | 2 | 0 | 0 | 3 |
| Anterior | 15 | 1 | 2 | 0 | 0 | 3 |
| Anterior | 16 | 1 | 1 | 3 | 0 | 5 |
| Anterior | 16 | 1 | 1 | 2 | 5 | 9 |
| Anterior | 16 | 1 | 1 | 1 | 9 | 12 |
| Anterior | 16 | 1 | 1 | 2 | 5 | 9 |
| Middle | 15 | 1 | 1 | 1 | 0 | 3 |
| Middle | 15 | 1 | 1 | 2 | 0 | 4 |
| Posterior | 16 | 1 | 2 | 1 | 7 | 11 |
| Posterior | 14 | 1 | 6 | 3 | 2 | 12 |
| Posterior | 15 | 1 | 5 | 3 | 3 | 12 |
| Posterior | 15 | 1 | 4 | 1 | 7 | 13 |
| Posterior | 15 | 1 | 2 | 4 | 2 | 9 |
| Posterior | 15 | 1 | 1 | 1 | 0 | 3 |

A defining feature of type II neuroblasts is their ability to make INPs which undergo a molecularly asymmetric cell division to self-renew and generate a GMC (Bello et al., 2008; Boone and Doe, 2008; Bowman et al., 2008; Izergina et al., 2009). Here we determine if embryonic INPs undergo asymmetric cell division. To identify INPs and their progeny, we used the INP marker *R9D11-tdTomato* (henceforth *9D11-tom*) (Bayraktar and Doe, 2013), and confirmed that it is expressed in embryonic INPs (Figs 3A,B). We also detected a deep ventral cluster of unrelated cells that expressed *9D11-tom* but not Dpn, but these can be excluded from analysis due to their distinct position (Fig. 3A, asterisk). Using this marker, we found that *9D11-tom*^+^ Dpn^+^ embryonic INPs undergo asymmetric cell division: they partition aPKC and Miranda to opposite cortical domains (Fig. 3C). To confirm that these GMCs generate post-mitotic neurons during embryogenesis, we stained for the neuronal marker Elav, and found that *9D11-tom* clusters contained Elav^+^ neurons (Fig. 3D). Additionally, axon fascicles from single type II neuroblast lineage clones were visible during embryogenesis (data not shown), confirming the production of embryonic-born neurons from type II lineages. We conclude that embryonic type II neuroblasts generate asymmetrically dividing INPs that produce GMCs and neurons during embryogenesis.

**Figure 3.**
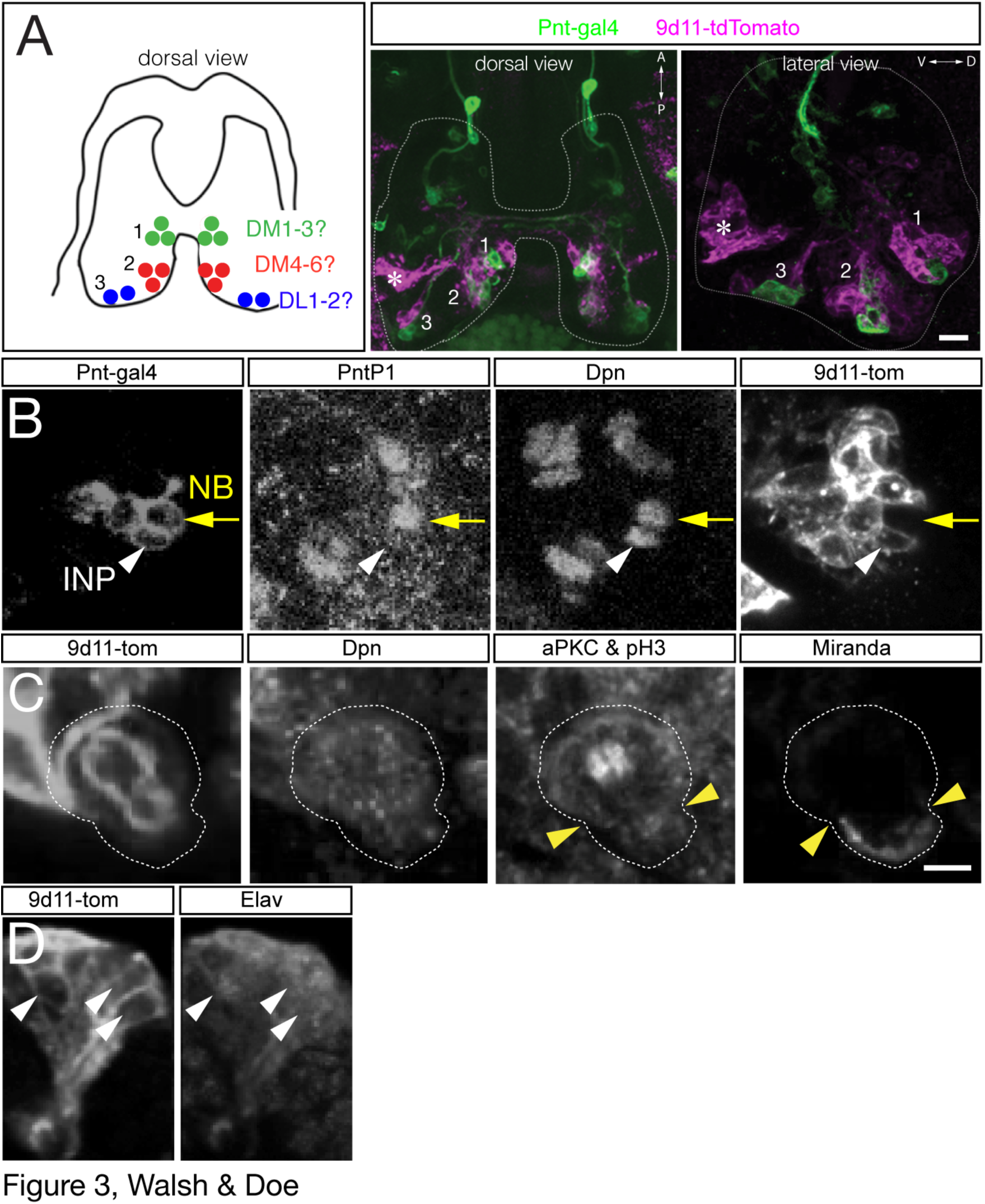
Embryonic INPs undergo asymmetric cell division. (**A**,**B**) *R9D11-tdTomato* (*9Dll-tom*) labels embryonic INPs and their progeny, but not type II neuroblasts. (A) Left: summary of type II neuroblast positions (dorsal view). Center and left panels: dorsal or lateral view of the three type II neuroblast clusters labeled with *Pnt-gal4* (green; type II neuroblasts and progeny) and *9D11-tom* (magenta; INPs and progeny). Note there is *9Dll-tom* expression at a deep ventral location that is not near any type II lineage (asterisk). Scale bar, 15 μm. (B) Type II neuroblast (*Pnt-gal4*^+^ PntP1^+^ Dpn^+^ *9Dll-tom*^+^ (yellow arrow); INP (*Pnt-gal4*^+^ PntPl^-^ Dpn^+^ *9D11-tom*^+^ (white arrowhead) at stage 16. Scale bar, 10 μm. **(C)** Embryonic INPs undergo asymmetric cell division. INPs were identified as *9D11-tom*^+^ Dpn^+^ and positioned within the middle cluster of neuroblasts in the dorsal posterior medial brain lobe. aPKC and pH3 are co-stained: aPKC is localized to the larger apical cell cortex (white cortex above arrowheads; future INP daughter cell) while pH3 decorates the mitotic chromosomes in the middle of the INP. Miranda is localized to the smaller basal cell cortex (cyan cortex below arrowheads; future GMC daughter cell). Scale bar, 5 μm. **(D)** Embryonic INPs generate embryonic-born neurons. Lateral view of a *9D11-tom*^+^ cluster in a stage 16 embryo. The post-mitotic neuronal marker Elav is detected in a subset of the *9D11-tom*^+^ cluster (white arrowheads), and axon projections can be observed (bottom left). Scale bar, 5 μm.

### Embryonic type II neuroblasts and INPs undergo quiescence

Type I central brain and thoracic neuroblast have been shown to undergo quiescence at the embryo-larval transition (Truman and Bate, 1988). Type II neuroblasts also undergo quiescence, because only the four mushroom body neuroblasts and a single lateral neuroblast maintain proliferation during the embryo-larval transition (Egger et al., 2008). In contrast, nothing is known about whether INPs undergo quiescence. To address this question, we counted the total number of INPs over time, as well as the number of mitotic INPs. We identified INPs as *9D11-tom*^+^ Dpn^+^ and mitotic INPs by immunoreactivity for phospho-histone H3 (pH3). We quantified INPs in each cluster independently as well as all INPs in each brain lobe (Fig. 4A). We observed a fairly constant number of INPs in each cluster from embryonic stage 14 to stage 17 (Fig. 4B), yet the number of proliferating INPs declined significantly over time, reaching zero by stage 17 (Fig. 4C). We conclude that the INPs enter quiescence by embryonic stage 17.

**Figure 4.**
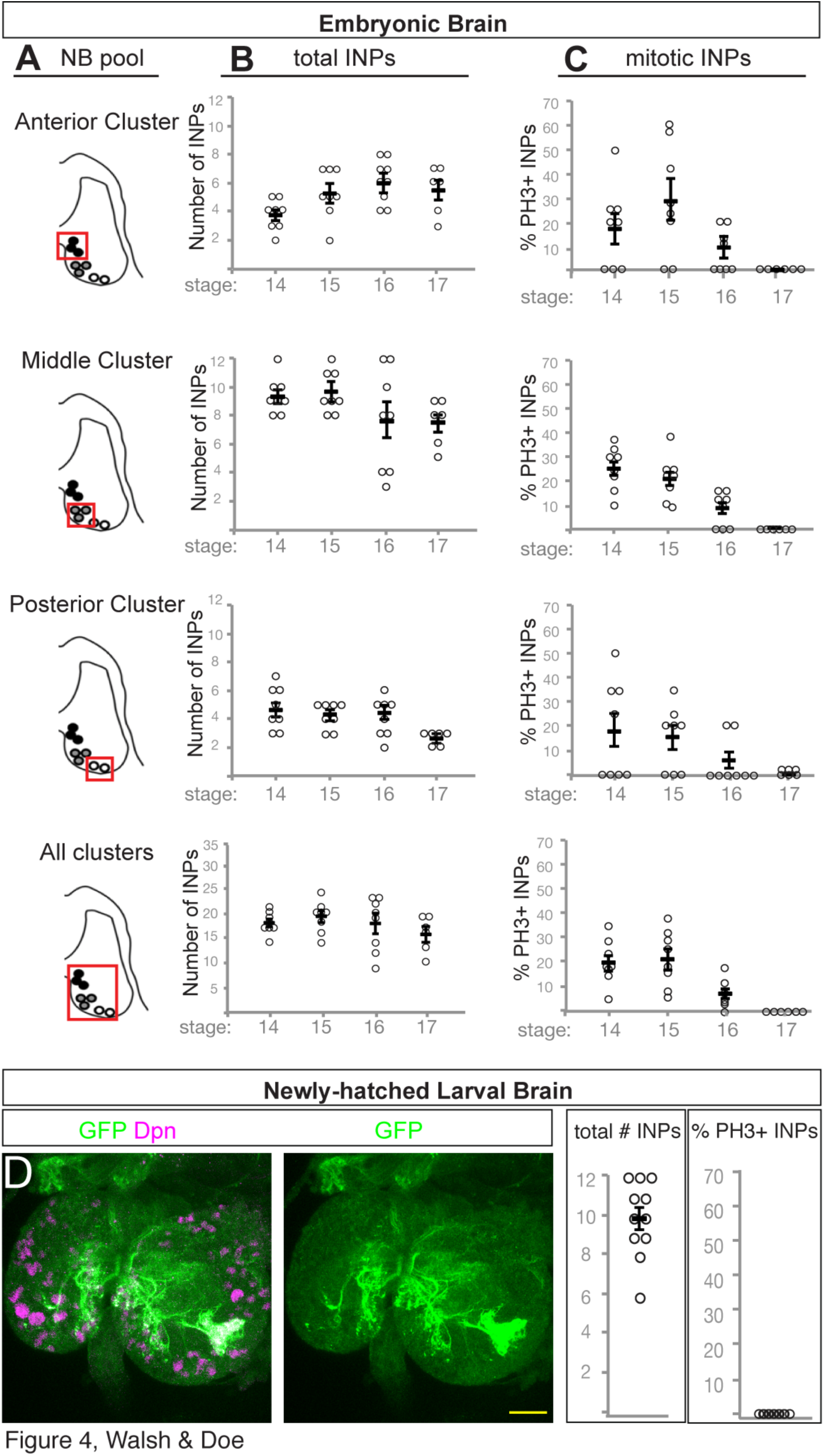
INPs undergo quiescence across the embryo-larval transition. **(A)** Schematic outlining the three pools of type II neuroblast INP progeny assayed in graphs to the right (red box). **(B)** Total number of INPs per pool at the indicated stages; INPs identified as 9D11-tom^+^ Dpn^+^ cells. **(C)** Number of phospho-histoneH3 (pH3)-positive mitotic INPs per pool at the indicated stages; INPs are identified as *9D11-tom*^+^ Dpn^+^ cells. Each circle represents the number of INPs in the cluster of neuroblasts shown in A; black bar represents the average, shown with SEM. **(D)** Quiescent INPs are present in the newly hatched larva. INPs marked with *9D11-gal4 UAS-tdTomato* (green); brain neuroblasts and INPs marked with Dpn (magenta). Anterior up, dorsal midline, dashed. Scale bar = 15 μm.

If INPs enter quiescence in the late embryo, we should be able to detect them in the newly hatched larvae, prior to production of larval born INPs made from type II neuroblasts that have re-entered the cell cycle. We assayed 0-4h newly-hatched larvae for Dpn and *9D11-tom* to mark the small quiescent INPs (Fig. 4D). We observed an average of 10 ± 2 *9D11-tom*^+^ Dpn^+^ cells in each brain lobe, and none of these INPs were mitotic (n=11; Fig. 4D). We conclude that INPs undergo quiescence in the late embryo and can persist into the larvae. The fate of these quiescent INPs – whether they resume proliferation, differentiate, or die – remains to be determined.

### Embryonic type II neuroblasts undergo a late temporal transcription factor cascade

Embryonic type I neuroblasts undergo a well-characterized temporal transcription factor cascade that generates GMC diversity and ultimately neuronal diversity. Most type I neuroblasts sequentially express Hunchback > Krüppel > Pdm > Cas > Grh (Kohwi and Doe, 2013), although late-forming neuroblasts can skip some of the early factors: neuroblast 3-3 begins the series with Krüppel (Tsuji et al., 2008) and NB6-1 begins the series with Cas (Cui and Doe, 1992). Due to the fact that type II neuroblasts are among the latest to form, it raises the possibility that they do not express any known temporal transcription factors.

We stained embryos for type II neuroblast markers (Dpn^+^ Ase^−^) and individual temporal identity transcription factors. We did not observe the first two temporal transcription factors, Hunchback or Krüppel, in any type II neuroblasts at any stage of development (data not shown). We next focused on the first type II neuroblast to form, which can be uniquely identified at late stage 11 (see Fig. 1). This early-forming neuroblast showed the temporal cascade of Pdm > Pdm/Cas > Cas > Cas/Grh > Grh (Fig. 5). All later-forming type II neuroblasts exhibited a more truncated temporal cascade of Cas > Cas/Grh > Grh (Fig. 5). We conclude that embryonic type II neuroblasts undergo a late temporal transcription factor cascade.

**Figure 5.**
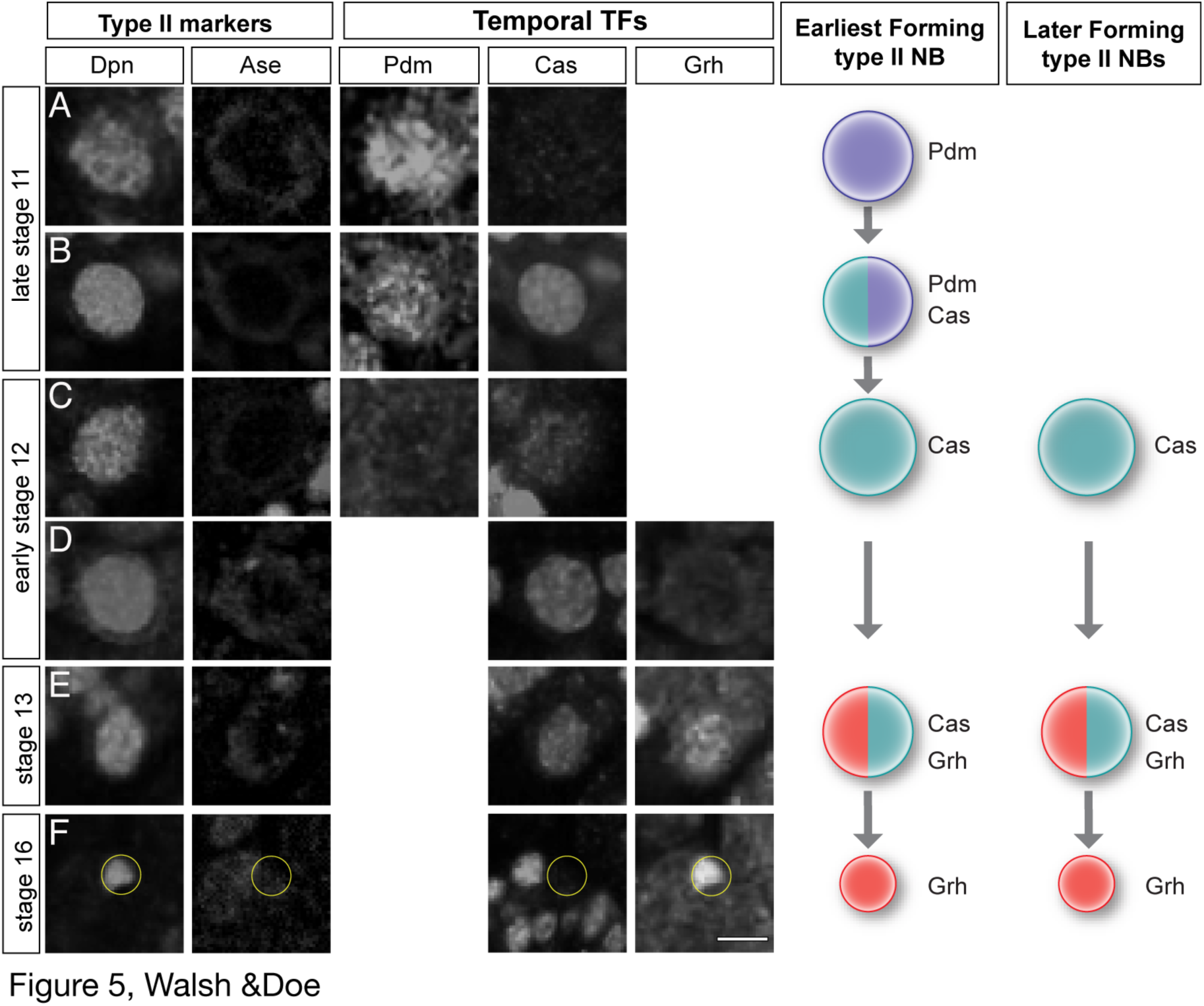
Embryonic type II neuroblasts express late temporal transcription factors. (A-F) Temporal transcription factor expression in the earliest type II neuroblast to form (posterior-most, see Fig. 1). Type II neuroblasts identified as Dpn^+^ Ase^-^ (left columns); temporal transcription factor expression reveals sequential expression of Pdm^+^ > Pdm^+^ Cas^+^ > Cas^+^ > Cas^+^ Grh^+^ > Grh^+^. Summarized at left; later-forming type II neuroblasts start the cascade with Cas. Scale bar = 10 μm.

### Embryonic INPs undergo a truncated temporal transcription factor cascade

Larval INPs undergo a temporal transcription factor cascade of Dichaete-Grh-Eyeless over their ~12 hour lifespan (Bayraktar and Doe, 2013). We wondered if the shorter timeframe of embryogenesis may result in shorter temporal transcription factor expression windows, a truncated temporal cascade, or perhaps a lack of all temporal transcription factor expression.

To identify embryonic INPs expressing known INP temporal transcription factors, we generated FLP-out clones using a heat shock FLP in mid-embryogenesis (4h-9h) and assayed brains containing a single type II neuroblast clone. We stained embryos for the clone marker, Dpn, and Ase to identify the neuroblast (Dpn^+^ Ase^-^) and INPs (Dpn^+^ Ase^+^), and one of the larval INP temporal transcription factors (Dichaete, Grh or Eyeless). We found that the early temporal factor Dichaete was detected in all INPs within the anterior and middle clusters (n=15 clones, anterior; n=12 clones, middle) (Fig. 6A,B; quantified in Table 2), but the posterior cluster contained no Dichaete^+ INPs^ at any stage (n=9 clones) (Fig. 6C; quantified in Table 2). The middle temporal factor, Grh, was only detected in a single INP next to Grh^+^ neuroblasts, but not next to Grh^-^ neuroblasts, suggesting that it is transiently inherited from the parental neuroblast, as is also observed in larval INP lineages (Bayraktar and Doe, 2013); we never detected Grh in INPs distant from the neuroblasts, as would be expected for a middle temporal transcription factor (data not shown). The late temporal factor Eyeless was never detected in INPs during embryogenesis (data not shown). We conclude that embryonic INPs undergo a temporal cascade that is truncated during the Dichaete window by entry into quiescence (Fig. 6E). It would be interesting to determine whether embryonic-born INPs express the later temporal factors Grh and Eyeless in the larvae, if they re-enter the cell cycle.

**Figure 6.**
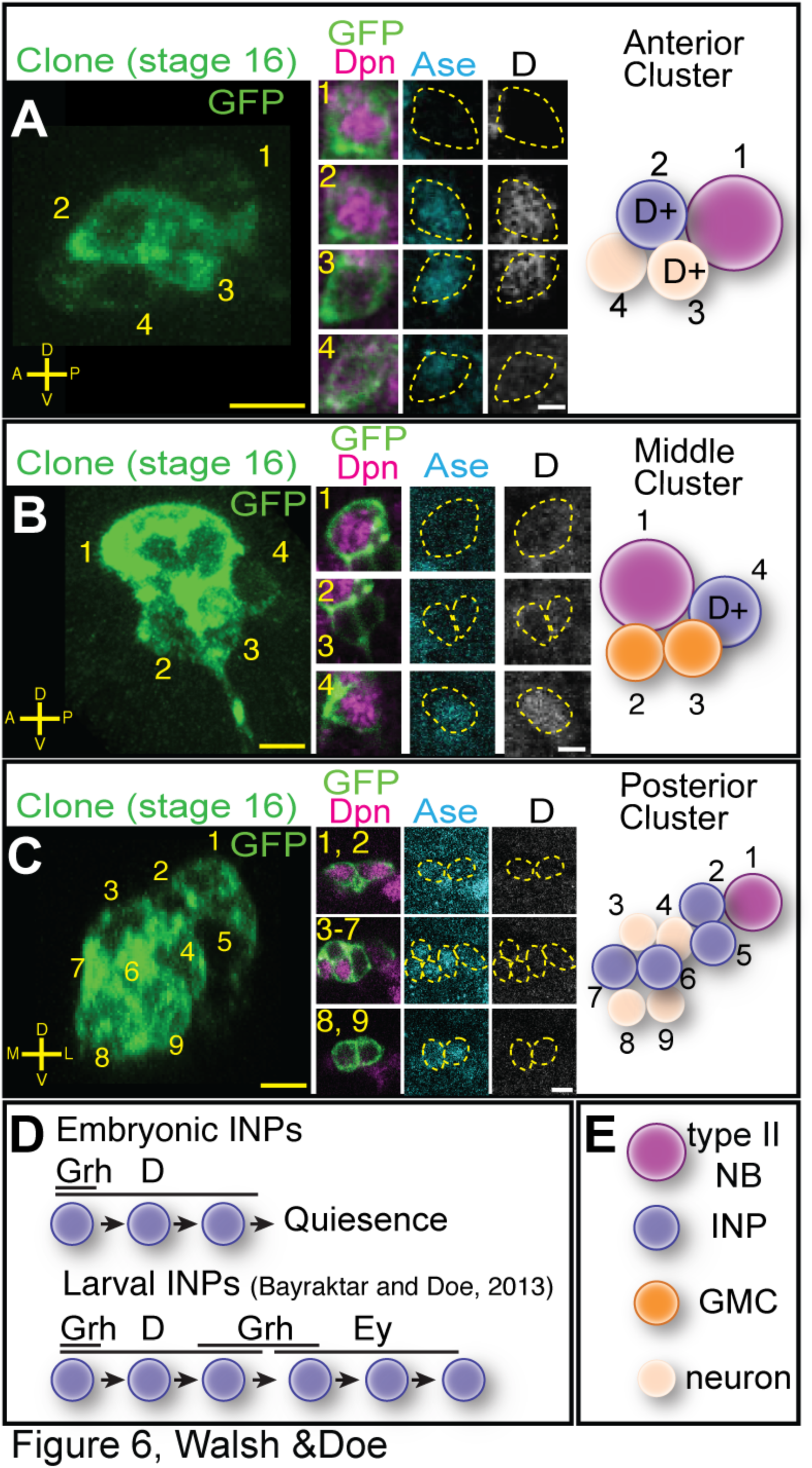
Embryonic INPs express the Dichaete temporal transcription factor. **(A)** Anterior cluster clone containing Dichaete (D)^+^ INPs. Four cell FLP-out clone at stage 16 (left) stained for the clone marker (GFP, green), Dpn (magenta), Ase (cyan) and D (white). The clone contains a type II neuroblast (1), a D^+^ INP (2) and two GMCs, one D^+^ and one D^-^ (3,4) **(B)** Anterior cluster clone containing D+ INPs. Four cell FLP-out clone at stage 16 stained the same as in (A) containing a type II neuroblast (1), one D^+^ INP (4), and two D^-^ GMCs (2,3). **(C)** Posterior cluster clone lacking D+ INPs. Nine cell FLP-out clone at stage 16 (left) stained the same as in (A) containing a type II neuroblast (1), four D^-^ INPs (2,5-7) and four D^-^ neurons (3,4,8,9). Scale bar 7 μm in clonal projections, 5 μm in insets. **(D)** Model for INP temporal factor expression; top, embryonic INPs from anterior and middle clusters; bottom, larval INP temporal factor expression (Bayraktar and Doe, 2013). **(E)** Cell type key for panels above.

**Table 2.** Dichaete is expressed in embryonic INPs. Each row represents a single neuroblast clone that is spatially separate from other clones in the embryonic brain. Stage, time of clone analysis.

| Cluster | Stage | Type II NB<br>*Dpn+ Ase-* | INP<br>*Dpn+Ase+* | D+ INP<br>*Dpn-Ase+ D+* |
| --- | --- | --- | --- | --- |
| Anterior | 14 | 1 | 1 | 1 |
| Anterior | 15 | 1 | 2 | 2 |
| Anterior | 15 | 1 | 1 | 1 |
| Anterior | 15 | 1 | 2 | 2 |
| Anterior | 16 | 1 | 1 | 1 |
| Anterior | 16 | 1 | 2 | 2 |
| Middle | 15 | 1 | 1 | 1 |
| Middle | 15 | 1 | 1 | 1 |
| Middle | 15 | 1 | 2 | 2 |
| Middle | 16 | 1 | 2 | 2 |
| Middle | 16 | 1 | 2 | 2 |
| Middle | 16 | 1 | 1 | 1 |
| Middle | 16 | 1 | 1 | 1 |

### Embryonic-born INPs contribute to the adult central complex

Embryonic type II neuroblasts produce neurons with contralateral projections, where they have been proposed to pioneer the fan shaped body neuropil of the central complex (Riebli et al., 2013). To determine if embryonic-born INP progeny persist into adulthood we used the FLEX-AMP system (Bertet et al., 2014) to permanently mark embryonic INPs and their progeny and trace them into the adult brain. FLEX-AMP uses a brief inactivation of temperature-sensitive Gal80 protein (by shifting to 29^o^C) to allow transient expression of Gal4, which induces FLP expression and the permanent expression of *actin-LexA LexAop-myr:GFP* (Fig. 7A). We crossed *R9D11-gal4* (expressed in embryonic INPs) to the FLEX-AMP stock and raised the flies at 18^o^C (negative control), 29^o^C (positive control), or with a 10 hour pulse of 29^o^C at late embryogenesis followed by 18^o^C for the rest of the fly’s life (“immortalization of embryonic progeny” experiment).

**Figure 7.**
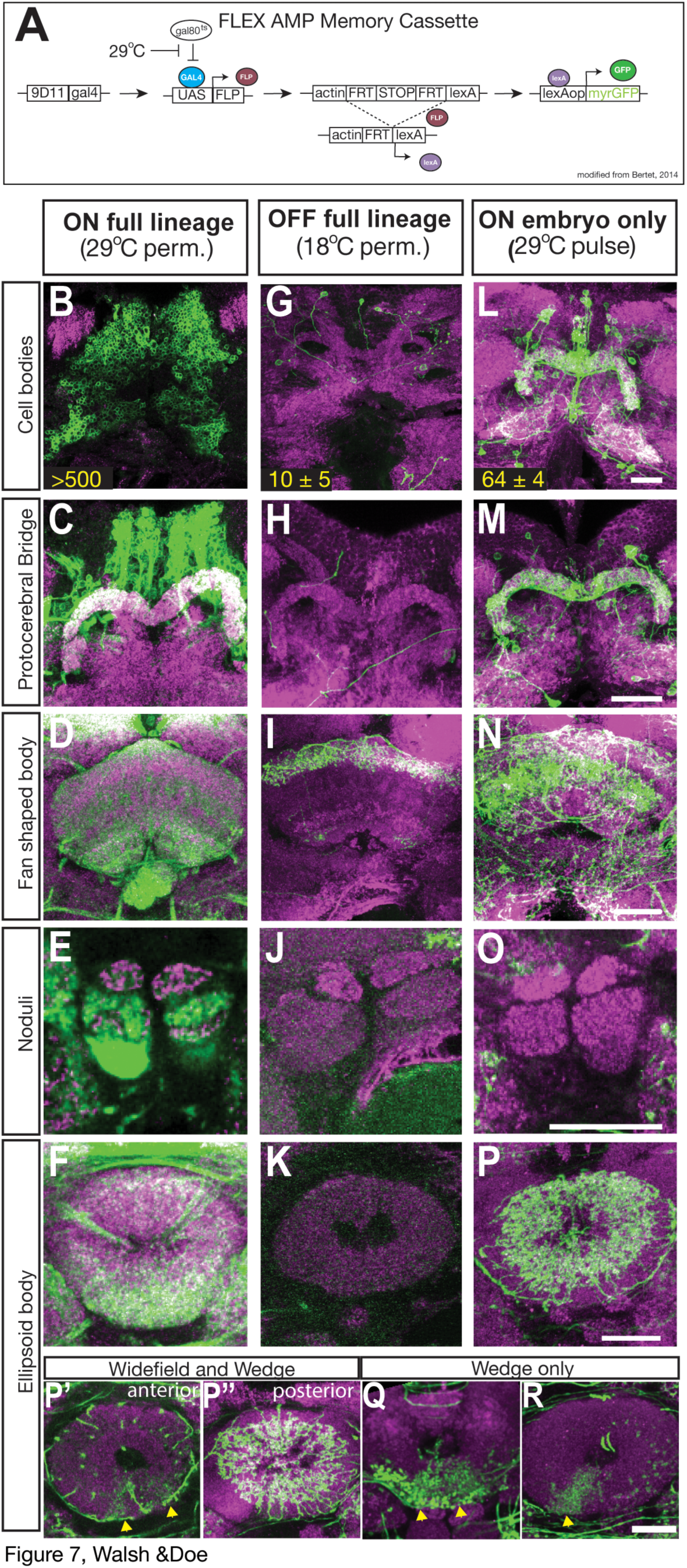
Embryonic INP progeny contribute to the adult central complex. **(A)** The FLEX AMP memory cassette used for immortalization of embryonic INPs into the adult brain; modified from Bertet et al. 2014. **(B-P)** Central complex neuropil regions from flies containing FLEX AMP memory cassette reared at different temperature regimes to permanently label neurons born within all development (29^o^C positive control), no stage of development (18^o^C negative control) or specifically during late embryogenesis (29^o^C pulse) stained for GFP (green) and NC82 (magenta). **(B-F)** Positive controls reared at 29^o^C from embryo to adult with over 500 (n= 4) of immortalized neurons innervating the PB, FB, EB and NO. **(G-K)** Negative control adult brains of flies reared at 18^o^C from embryo to adult showing 10 ± 5 (n=5) neurons from the adult *9D11-gal4* pattern innervating only the dorsal region of the FB. **(L-R)** Experimental adult brains from flies reared for 6 hour pulse at 29^o^C at late embryonic stages, then reared at 18^o^C until adult (see methods); there are 64 ±4 (n=12) neurons that innervate the PB, FB, EB but not the NO. **(P’-R)** Experimental adult brains with differences in innervation pattern within the EB (n=12). **(P’)** Single z plane from anterior region shown in (P) with innervation of two wedges within the EB (yellow arrows) seen within 12/12 brains. **(P”)** Single z plane from posterior region shown in (P) showing wide field neuron innervation within the EB seen within 9/12 brains. **(Q)** EB with innervation of two wedges, lacking the wide field innervation (n=1). **(R)** EB with innervation of one wedge, lacking the wide field innervation (n=1). Abbreviations: PB (protocerebral bridge), FB (fan shaped body), EB (ellipsoid body), NO (noduli). Scale bar = 20μm.

We found robust labeling of >500 neurons in the positive control brains raised at 29^o^C, including many cell bodies innervating the protocerebral bridge, fan shaped body, ellipsoid body and noduli (Fig. 7B-H). The negative control (18^o^C permanently) showed labeling of just ~10 neurons that project to the dorsal part of the fan shaped body (Fig. 7G-K), which is similar to the adult pattern of R9D11 (FlyLight). We suspect the “leaky” expression at 18 ^o^C may reflect the inefficiency of Gal80 repression in these adult neurons. Importantly, FLEX-AMP immortalization of embryonic INP progeny showed labeling of additional neurons (64 ± 4) that project to three central complex regions: the protocerebral bridge, a large portion of the fan shaped body and the ellipsoid body, but notably not the noduli (Fig. 7 L-P). Within the ellipsoid body, we observed variation in labeling. Most brains contained one to two wedge neurons (arrows in Fig. 7P’) and widefield neuron innervation throughout the posterior region of the ellipsoid body (Fig. 7P”, n= 12). Interestingly, a few brains contained only the wedge neurons suggesting the widefield neuron innervation may be an early-born neuron within the lineages (See Discussion) (n= 3/12, Fig. 7 Q, R). Additionally, FLEX-AMP immortalization of embryonic INP progeny identified neurons innervating the central complex accessory neuropils lateral accessory lobe (LAL) and the Gall, which were never labeled in the 18^o^C negative control (Fig. S1). We conclude that embryonic INPs generate progeny that persist into the adult brain, and innervate three neuropils of the central complex.

## DISCUSSION

It has been difficult to link embryonic neuroblasts to their larval counterparts in the brain and thoracic segments due to the period of quiescence at the embryo-larval transition, and due to dramatic morphological changes of the CNS that occur at late embryogenesis. Recent work has revealed the embryonic origin of some larval neuroblasts: the four mushroom body neuroblasts in the central brain and about twenty neuroblasts in thoracic segments (Kunz et al., 2012; Lacin and Truman, 2016). Here we use molecular markers and clonal analysis to identify all eight known type II neuroblasts in each brain lobe and show they all form during embryogenesis, perhaps the last-born central brain neuroblasts. We were unable to individually identify each neuroblast, however, due to their tight clustering, movements of the brain lobes, and lack of markers for specific type II neuroblasts.

The single previously reported embryonic type II neuroblast formed from PntP1^+^ neuroectodermal cells with apical constrictions called a placode (Hwang and Rulifson, 2011). We have not investigated this neuroectodermal origin of type II neuroblasts in much detail, but we also observe multiple type II neuroblasts developing from PntP1^+^ neuroectoderm (data not shown). In the future, it would be interesting to determine whether all type II neuroblasts arise from PntP1^+^ neuroectoderm or from neuroectodermal placodes. Interestingly, one distinguishing molecular attribute of type II neuroblasts is PntP1, which is not detected in type I neuroblasts (Zhu et al., 2011; Xie et al., 2016). Thus, a candidate for distinguishing type I / type II neuroblast identity is EGF signaling, which can be detected in the three head placodes (de Velasco et al., 2007; Hwang and Rulifson, 2011) and is required for PntP1 expression (Gabay et al., 1996). Clearly there are more PntP1+ neuroectodermal cells than there are type II neuroblasts, however, which may require expression of an EGF negative regulator such as Argos (Rebay, 2002) to divert some of these neuroectodermal cells away from type II neuroblast specification. The earliest steps of type II neuroblast formation represent an interesting spatial patterning question for future studies.

Now that we have identified the embryonic type II neuroblasts, it is worth considering whether there are differences between embryonic and larval type II neuroblasts or their INP progeny. To date, molecular markers do not reveal any differences between embryonic and larval type II neuroblasts, with the exception that embryonic neuroblasts transiently express the temporal transcription factor Pdm (see below). Are there differences between embryonic and larval INPs? Larval INPs mature over a period of six hours and then divide four to six times with a cell cycle of about one hour (Bello et al., 2008). In contrast, embryonic INPs may have a more rapid maturation because we see Elav^+^ neurons within 9D11^+^ INP lineages by stage 14, just 3 hours after the first type II neuroblast forms. We found that INPs undergo quiescence at the embryo-larval transition, as shown by the pools of INPs at stage 16 that do not stain for the mitotic marker pH3. The fate of these quiescent INPs – whether they resume proliferation, differentiate, or die – remains to be determined.

Neuroblasts in the embryonic VNC use the temporal transcription factor cascade Hunchback (Hb) > Krüppel > Pdm > Cas > Grh to generate neural diversity (Brody and Odenwald 2002; Kohwi et al., 2013; Allan and Thor, 2015; Kang and Reichert, 2015; Doe, 2017). Here we show that the type II neuroblasts are among the last neuroblasts to form in the embryonic brain, and that they sequentially express only the late temporal transcription factors Pdm (in the earliest-forming neuroblast) followed by Cas and Grh (in all eight type II neuroblasts). It is unknown why most type II neuroblasts skip the early Hb > Kr > Pdm temporal transcription factors; perhaps it is due to their late time of formation, although several earlier-forming thoracic neuroblasts also skip Hb (NB3-3), Hb > Kr (NB5-5), or Hb > Kr > Pdm (NB6-1) (Cui and Doe, 1992; Tsuji et al., 2008; Benito-Sipos et al., 2010). This is another interesting spatial patterning question for the future.

Type I neuroblasts show persistent expression of the temporal transcription factors within neurons born during each window of expression (i.e. a Hb^+^ neuroblast divides to produce a Hb^+^ GMC which makes Hb^+^ neurons). In contrast, we find that type II neuroblasts do not show persistent Cas or Grh expression in INPs born during each expression window (data not shown). Both transcription factors can be seen in INPs immediately adjacent to the parental neuroblast, but not those more distant (data not shown). This shows that Cas and Grh are down regulated in INPs rather than maintained in the INP throughout its lineage and into all its post-mitotic neural progeny. The function of Pdm, Cas and Grh in embryonic type II neuroblasts awaits identification of specific markers for neural progeny born during each expression window.

During larval neurogenesis, virtually all INPs sequentially express the temporal transcription factors Dichaete > Grh > Eyeless (Bayraktar and Doe, 2013). In contrast, embryonic INPs express only Dichaete. These data, together with the short time frame of embryogenesis, suggests that INP quiescence occurs during the Dichaete window, preventing expression of the later Grh > Ey cascade. Interestingly, INPs in the posterior cluster completely lack Dichaete, suggesting they may be using a different temporal transcription factor cascade. The posterior cluster type II neuroblasts are likely to be the DL1-DL2 type II neuroblasts, which have never been assayed for the Dichaete > Grh > Eyeless cascade in larval stages. Perhaps these two neuroblasts use a novel temporal cascade in both embryonic and larval stages.

Larval type II neuroblasts produce many intrinsic neurons of the adult central complex (Bayraktar and Doe, 2013; Ito et al., 2013; Yu et al., 2013). Here we show that embryonic INPs also produce neurons that contribute to the adult central complex. Our data show ~54 neurons (64 minus the 10 due to “leaky” expression) born from embryonic-born INPs survive to adulthood and innervate the central complex. It is likely that this is an underestimate, however, because (1) 9D11-gal4 expression is lacking from a few INPs in the embryonic brain and (2) the time to achieve sufficient FLP protein levels to achieve immortalization may miss the earliest born neurons. The variation in immortalization of the wide field ellipsoid body neuron may represent a neuron born early in the type II lineages, thus unlabeled in a subset of embryos. Additionally, some embryonic born neurons may perform important functions in the larval/pupal stages but die prior to eclosion.

Further studies will be required to understand the function of neurons born from embryonic type II lineages. It remains to be experimentally determined whether some or all embryonic progeny of type II neuroblasts (a) remain functionally immature in both the larval and adult brain, but serve as pioneer neurons to guide larval-born neurons to establish the central complex, (b) remain functionally immature in the larval brain, but differentiate and function in the adult central complex, or (c) differentiate and perform a function in both the larval and adult CNS. It will be informative to selectively ablate embryonic-born neurons and determine the effect on the assembly of the larval or adult central complex, and their role in generating larval and adult behavior.

## MATERIALS AND METHODS

### Fly stocks

The chromosomes and insertion sites of transgenes (if known) are shown next to genotypes. Unless indicated, lines were obtained from Bloomington stock center (FlyBase IDs shown). Enhancer gal4 lines and reporters: *P[GAL4]pnt^14-94^* (III) (gift of Jan Lab), *R9D11-gal4 (III*, *attP2*), *R9D11-CD4-tdTomato* (*III*, *attP2*), *10XUAS-IVS-mCD8::GFP* (*III*, *su(Hw)attP2*) (referred to as *UAS-GFP*). hs FLPG5;;MCFO (I and III; FBst0064086). For FLEXAMP experiment, *y*,*w*,*UAS-FLP*; *tubGAL80ts/CyO*; *R9D11-gal4/TM3* and *13Xlex-Aop2-myr::GFP*; *tubGAL80ts/CyO*; *P{nSyb(FRT.stop)LexA.p65}*.

### Immunofluorescent staining

Primary antibodies were rat anti-Dpn (1:50, Abcam; Eugene, OR, USA), guinea pig anti-Dpn (1:1000, Jim Skeath; Washington Univ.), chicken anti-GFP (1:1000, Aves Laboratories, Tigard, OR), guinea pig anti-D (1:500, John Nambu; Univ. Massachusetts, Amherst), rabbit anti-Ey (1:2500, Uwe Walldorf; Germany), rabbit anti-phospho-Histone H3 (ser 10) (1:20,000, Millipore, Temecula, CA), rabbit anti-PntP1 (1:1000, Jim Skeath; Washington Univ.), rat anti-Grh (1:1000, Stefan Thor), rabbit anti-DsRed (1:1000, Clontech Laboratories, Mountain View, CA, USA), rabbit anti-Ase (1:1000, Cheng-Yu Lee; Univ. Michigan), mouse anti-Hunchback (1:500; Abcam; Eugene, OR, USA), guinea pig anti Krüppel (1:500, Doe Lab), rat anti-Pdm2 (1:1000 Abcam; Eugene, OR, USA), guinea pig anti-Asense (1:1000; Hongyan Wang, NUS/Duke, Singapore), rabbit anti-Cas (1:1000, Ward Odenwald, distributed by the Doe lab), mouse anti-NC82 (1:200, Developmental Studies Hybridoma Bank). Secondary antibodies were from Molecular Probes (Eugene, OR, USA) or Jackson Immunoresearch (West Grove, PA, USA) used at 1:400.

Embryos were blocked overnight in 0.3% PBST (1X phosphate buffered saline with 0.3% Triton X-100) with 5% normal goat serum and 5% donkey serum (PDGS) (Vector Laboratories, Burlingame, CA, USA), followed by incubation in primary antibody overnight at 4^o^C. Next, embryos underwent four washes 15 minutes each in PDGS, followed by a 2 hour secondary antibody incubation at 25^o^C. After secondary, embryos were either dehydrated with ethanol and mounted in dibutyl phytalate in xylene (DPX) according to Janelia protocol (Wolff et al., 2015) or were cleared with a glycerol series: 25% for 10 minutes, 50% for ten minutes, 90% for ten minutes then into 90% glycerol with 4% n-propyl gallate overnight before imaging.

Larval brains were dissected in PBS, fixed in 4% formaldehyde in PBST for 25 min, rinsed 30 minutes PBST, and blocked in PDGS overnight at 4^o^C. Staining as above for embryos, but after secondary were mounted in Vectashield (Vector Laboratories, Burlingame, CA, USA).

Adult brains were fixed in 2% formaldehyde in PBST, rinsed, and blocked in PDGS with 0.5% Triton. Brains were incubated in primary antibodies for four days at 4^o^C, then in secondary antibodies for two days at 4 ^o^C. Brains were mounted in DPX according to Janelia protocol.

### Clones

For type II clones, *P[GAL4]pnt^14-94^* (III) x *hs FLPG5;;MCFO* (I and III; FBst0064086) embryos were collected for four hours at 25^o^C, aged four hours and heat shocked at 37^o^C for 12 minutes, then left to develop until desired stages.

### FLEX-AMP immortalization of embryonic INPs

The FLEXAMP experiment used 1-3 day old adult females from crossing: *y*,*w*, *UAS-FLP*; *tubGAL80ts/CyO*; *R9D11-gal4/TM3* to *13Xlex-Aop2-myr::GFP*; *tubGAL80ts/CyO*; *P{nSyb(FRT.stop)LexA.p65}* to permanently label embryonic INPs (Bertet et al., 2014). Negative controls were raised continuously at 18^o^C to maintain Gal80 repression; positive controls were raised continuously at 29^o^C inactivate Gal80 and allow 9D11-gal4 expression. To “immortalize” embryonic INPs and their progeny, we exposed embryos aged 5-6 hours to 29^o^C for ten hours to a allow *R9D11-gal4* expression and then shifted all unhatched embryos to 18^o^C to block *R9D11-gal4* expression during larval, pupal and adult stages.

### Cell proliferation analysis

Number of proliferating INPs was calculated by dividing the number pH3 positive by the number of total INPs within each cluster of neuroblast at different stages. Each circle represents one cluster of INPs. Error bars represent standard error of the mean.

### Imaging

Images were captured with a ZeissLSM700 or ZeissLSM710 confocal microscope with a z-resolution of 1.0 micron, and processed in the open source software FIJI (http://fiji.sc) and Photoshop CS5 (Adobe, San Jose, CA, USA). Figures were made in Illustrator CS5 (Adobe, San Jose, CA, USA). Three-dimensional brain reconstructions in Figs. 3 and 6 were generated using Imaris (Bitplane, Zurich, Switzerland).

## Acknowledgements

We thank Luis Sullivan, Emily Sales, Laurina Manning, and Sen-Lin Lai for fly stocks, technical assistance and helpful discussions; Sen-Lin Lai and Volker Hartenstein for comments on the manuscript; Jim Skeath, John Nambu, Uwe Walldorf, Stefan Thor, Cheng-Yu Lee, Claude Desplan, Hongyan Wang, and Ward Odenwald for reagents; and Fernando Diaz-Benjumea and Volker Hartenstein for sharing unpublished work. We acknowledge the Bloomington Drosophila Stock Center (NIH P40OD018537) and the Developmental Studies Hybridoma Bank (DSHB).

## Author contributions

KTW and CQD conceived the project; KTW performed all the experiments; KTW and CQD wrote the manuscript.

## List of Symbols and Abbreviations

Ase: (Asense)
Dpn: (Deadpan)
D: (Dichaete)
EB: (ellipsoid body)
FB: (fan shaped body)
GMC: (ganglion mother cell)
Grh: (Grainy head)
Hb: (Hunchback)
INPs: (intermediate neural progenitors)
Kr: (Krüppel)
NO: (noduli)
Pdm: (Nubbin/Pdm2)
PntP1: (Pointed P1)
PB: (protocerebral bridge)
R9D 11-tdTomato: (9D 11-tom)

---

**Supplementary Figure 1.**
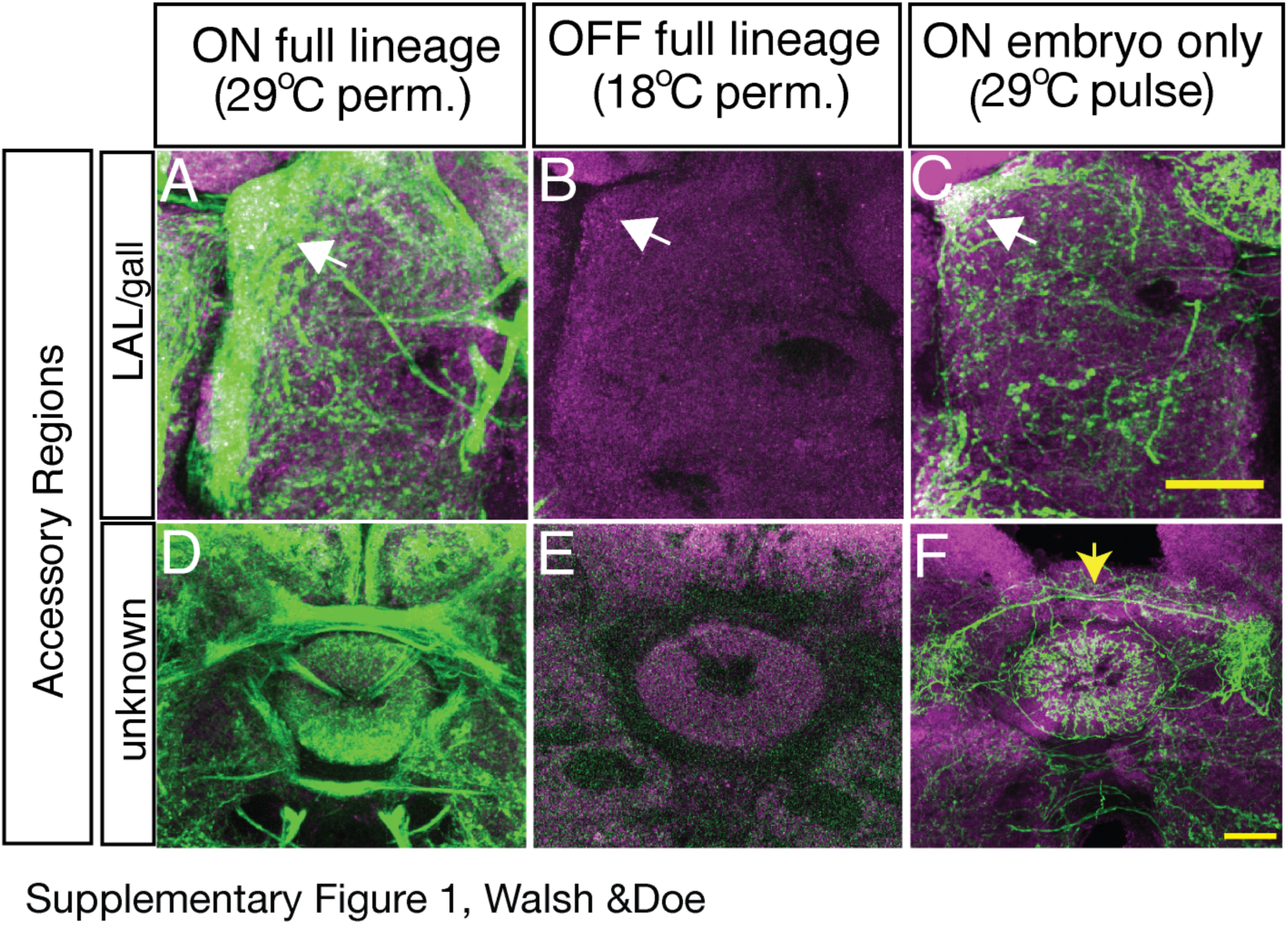
Embryonic INP progeny contribute to the adult Lateral Accessory Lobe (LAL) and Gall neuropils. **(A-F)** Staining of central complex accessory regions in FLEX AMP positive control (A, D), negative control (B, E) and embryo-only (C, F) groups. The LAL and gall (white arrows in top panel) are strongly innervated in positive control (A), negative in control (B), and diffusely innervated in embryonic labeled brain (note strong density within gall, arrow) (C). An unknown region adjacent to ellipsoid body is densely innervated in the positive control (D), absent in negative control (E), and innervated sparsely in the embryonic-only brain (F). Note the commissural axons within the pattern in (F, yellow arrow). Scale bar = 20 μm.

